# RIPK3 inhibition prevents KRAS mutant p53 deficient lung tumor progression by impeding MDSCs

**DOI:** 10.1101/853978

**Authors:** Asha Jayakumar

## Abstract

KRAS mutant p53 deficient (KP) non-small cell lung carcinoma (NSCLC) lacks targeted therapies. Existing treatments for lung cancer cause resistance and result in toxicities requiring novel effective therapies. By targeting mechanisms causing resistance such as myeloid derived suppressor cells (MDSCs), KP tumors could be inhibited. MDSCs are functionally diverse and suppress T cells in many cancers. RIPK3, a cell death inducing enzyme, also functions as a signaling component producing cytokines that mediate the suppressive function of MDSCs. Partial deletion of RIPK3 in myeloid cells reduced KP tumor growth. This also reduced accumulation of MDSCs with a consistent increase in antigen specific IFNγ producing CD8 T cells. Inhibiting RIPK3 with a small molecule inhibitor such as GSK 872 effectively reduces RIPK3 activity in myeloid cells including MDSCs and reduces growth of small and large KP tumors. GSK 872 in combination with checkpoint inhibitors such as anti PD-1 and anti CTLA-4 further decreased KP tumor size. Together, our findings show that inhibiting RIPK3 in MDSCs is effective in inhibiting KP NSCLC and is a viable therapeutic option for improving existing immunotherapeutic treatments.

## Introduction

Lung cancer is a major cause of death among people worldwide and in the United States (1). Non-small cell lung carcinoma (NSCLC) affects about 80% of lung cancer patients and lacks effective treatments. Among NSCLC, KRAS mutant p53 deficient (KP) lung cancer is prevalent among smokers and is a common cause of mortality (2). KP NSCLC is highly resistant to existing treatments indicating the need for new therapies. Several reports show that targeting tumor cells directly can reduce tumor progress. However, these methods lead to some form of resistance to treatment (2). One of the biggest challenges in NSCLC treatment is increased expression of checkpoint molecules that suppress the function of T cells (3). PD-1 and CTLA-4 are examples of checkpoint molecules that are expressed on immune cells in a tumor environment (4). Checkpoint blockade is a successful mode of immunotherapy that has been highly effective in NSCLC treatment (3). However, not all patients respond favorably to this treatment suggesting that alternative treatments are necessary for non-responders. Therefore, there is an intense need to develop treatments that improve the efficacy of immunotherapy with checkpoint inhibitors.

High immune cell infiltration in tumors show that they are hot tumors whereas cold tumors have low immune cell infiltration (5). Generally immune cell infiltration results in better response to therapy; however, increased expression of checkpoint molecules on immune cells reduces their efficacy. Other obstacles to treatment include increased frequency of suppressive immune cells such as myeloid derived suppressor cells (MDSCs) (6). MDSCs are generated from bone marrow cells by exposure to inflammatory cytokines and growth factors (7). They suppress T cells and also directly promote inflammation, tumor growth, and metastasis. A characteristic feature of MDSCs is their ability to multitask, thereby making it challenging to effectively reduce their tumor promoting function.

Receptor interacting protein kinases such as receptor interacting protein kinase 1 (RIPK1) and RIPK3 induce necroptotic cell death but also have an alternate function in signaling by inducing cytokines (8). We showed that targeting MDSCs by inhibiting RIPK3 signaling, changes the ability of MDSCs to promote Th17 cells in intestinal cancer (9). These results also showed that RIPK3 induced cytokines in MDSCs could support their suppressive function. The role of RIPK3 in NSCLC and suppressive function of MDSCs has not been studied. Therefore, we asked if inhibiting RIPK3 in NSCLC would inhibit the suppressive function of MDSCs and could effectively improve the generation of cytotoxic T cells. To answer this question, we used a murine tumor model of KP NSCLC expressing OVA antigen to show that a RIPK3 inhibitor GSK 872 could inhibit tumor growth. Using a RIPK3-GFP floxed mouse, we deleted RIPK3 in myeloid cells and showed that KP tumor growth was inhibited. These findings show that targeting RIPK3 could be a novel method to inhibit KRAS mutant NSCLC and improve checkpoint inhibitor treatment.

## Materials and methods

### Tumor models

C57BL/6J and OT-1 transgenic (Tg) mice were purchased from Charles River Laboratories. All mouse protocols were approved by the Yale University Institutional Animal Care and Use Committee. RIPK3 inhibitor GSK 872 (N-(6-(Isopropylsulfonyl)quinolin-4-yl)benzo[d]thiazol-5-amine) was purchased from Millipore Sigma and frozen stocks were maintained in DMSO. For KP tumors, 5 × 10^5^ KP cells were implanted subcutaneously in WT male C57BL/6J mice. At day 7 to 10 after tumor implantation, mice were treated every 3 days with GSK 872 at 0.5μg/gm body weight. Each mouse in the control group received 1μl DMSO per 200μl of PBS. Tolerability of GSK 872 was assessed by observing mice for lethargy or physical signs of allergic reaction at 24 hours after treatment with this inhibitor and every 3 days thereafter. Mice were treated with anti-PD-1 or anti-CTLA-4 antibodies at 100μg/mouse every week for 4 weeks. CD8 T cells were depleted by treating mice with anti CD8α antibody every week at 100ug/mouse. LysM Cre RIPK3-GFP floxed mice were maintained by breeding LysM-Cre and RIPK3 GFP floxed mice on C57BL/6J background. 5 × 10^5^ KP cells were implanted subcutaneously in male LysM-Cre RIPK3-GFP fl/WT mice. RIPK3 GFP floxed mice were kindly provided by Dr. Francis Chan at the University of Worcester. Genotype was confirmed by PCR analysis. Tumor volume was monitored every week and calculated with the following formula: short diameter^2^ x (long diameter)/2. KP tumor cells were regularly tested for Mycoplasma and found to be contamination-free.

### Adoptive cell therapy with OT-1 Tg CD8 T cells

Splenocytes from OT-1 Tg mice were treated with SIINFEKL peptide (0.75 μg/ml) for 5 to 7 days. T cell media included recombinant murine IL-2 at 10 units/ml. Media was replaced at day 3 and 5. Live cells specific for SIINFEKL peptide were collected by density gradient centrifugation using ficoll. 2 × 10^5^ cells were injected retro-orbitally per mouse at day 22 and 35 after KP tumor implantation.

### Flow cytometry

Splenocytes obtained by mechanical disruption were stained with CD45, CD4, CD8, TCRβ, CD11b, Gr-1, Ly6C, Ly6G, CD11c, and F4/80. MDSCs were identified using the gating strategy in Supplementary figure 1. For intracellular cytokine staining, splenocytes were activated with 5μg/ml of SIINFEKL peptide for 72 hours and brefeldin A was added during the last 4 hours of culture before fixation. Cells were surface stained, permeabilized with permeabilization buffer (eBioscience), and stained with anti-IFNγ. Viable CD45^+^ cells were used for all analysis.

### Statistical analysis

Data are presented as means ± SEM. Students t-test were used to determine statistical significance. p value less than 0.05 was considered significant.

## Results and Discussion

### RIPK3 in MDSCs promotes KP tumors

RIPK3 dependent ERK1/2 is a novel signaling pathway that accompanies pathogenic cell death or necroptosis that has been identified as a target for treating inflammatory diseases (8). RIPK3 signaling induces inflammatory cytokines in myeloid cells such as macrophages and MDSCs (9,10). We showed that RIPK3 signaling in MDSCs produces cytokines required to generate Th17 cells. In MDSCs, RIPK3 signaling also produces COX-2 (data not shown) and IL-1β, which are both required for the T cell suppressive function of MDSCs (7). This suggests that RIPK3 signaling in MDSCs could also suppress CD8 T cells to allow tumor growth. Therefore, we reasoned that inhibiting RIPK3 signaling in MDSCs could inhibit its suppressive function and reduce tumor growth. To test this rationale, we bred RIPK3-GFP floxed mice with LysM Cre mice to delete RIPK3 in myeloid cells. In RIPK3-GFP floxed mice, the RIPK homotypic interaction motif (RHIM) domain is linked to a GFP tag and deletion of this domain in Cre expressing cells deletes RIPK3 signaling (11). LPS induced signaling results in activation of RIPK3 RHIM to induce the signaling function of RIPK3 (8). In tumor bearing mice, MDSCs are most abundant among myeloid cells indicating that in LysM Cre RIPK3-GFP floxed mice, MDSCs would lack RIPK3. Heterozygous deletion of RIPK3 in LysM Cre mice, reduced RIPK3-GFP expression in CD11b+ cells and monocytes but not in neutrophils, DCs, and non-myeloid cells (CD11b-cells) (Supplementary fig. 1). This suggests that LysM Cre RIPK3 fl/wt mice with tumors would have a predominantly MDSC specific deletion of RIPK3 as MDSCs are similar to monocytes and neutrophils in WT mice. Additionally, reduction of RIPK3-GFP expression in MDSCs is consistent with the deletion of RIPK3-GFP in CD11c+ DCs as shown previously (11). The volume of KP tumors were reduced in LysM Cre RIPK3 fl/wt mice compared to WT mice (Fig. 1), showing that RIPK3 in MDSCs is important for the progression of KP tumors. CD11b+ myeloid cells and M-MDSCs were reduced in LysM Cre RIPK3 fl/wt mice (Fig. 2A).

**Figure 1.**
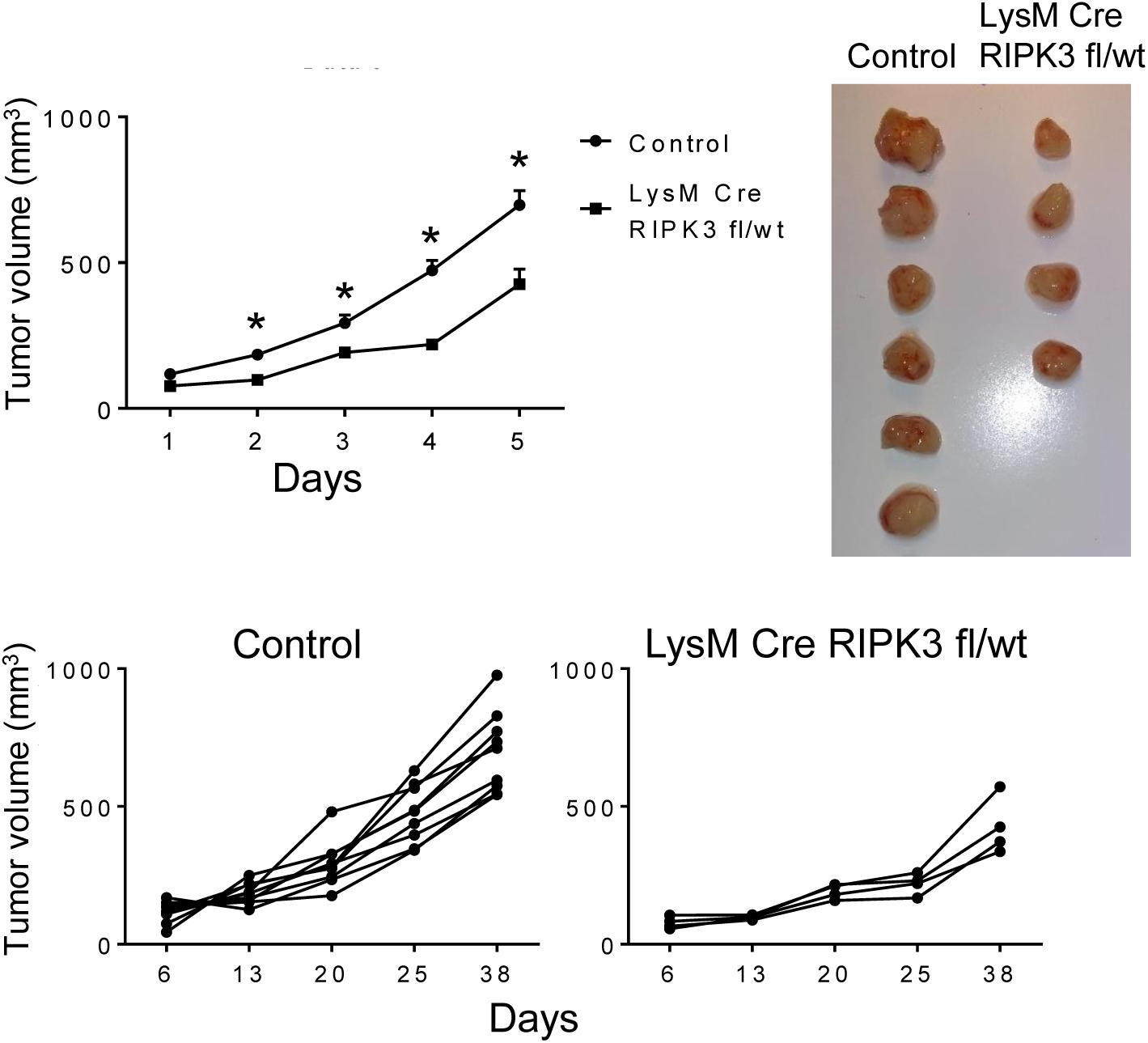
RIPK3 in myeloid cells promote KP lung tumors. KP lung tumor growth in WT and LysM Cre RIPK3-GFP fl/wt mice (n = 4 to 8 mice/group), * p< 0.05.

**Figure 2.**
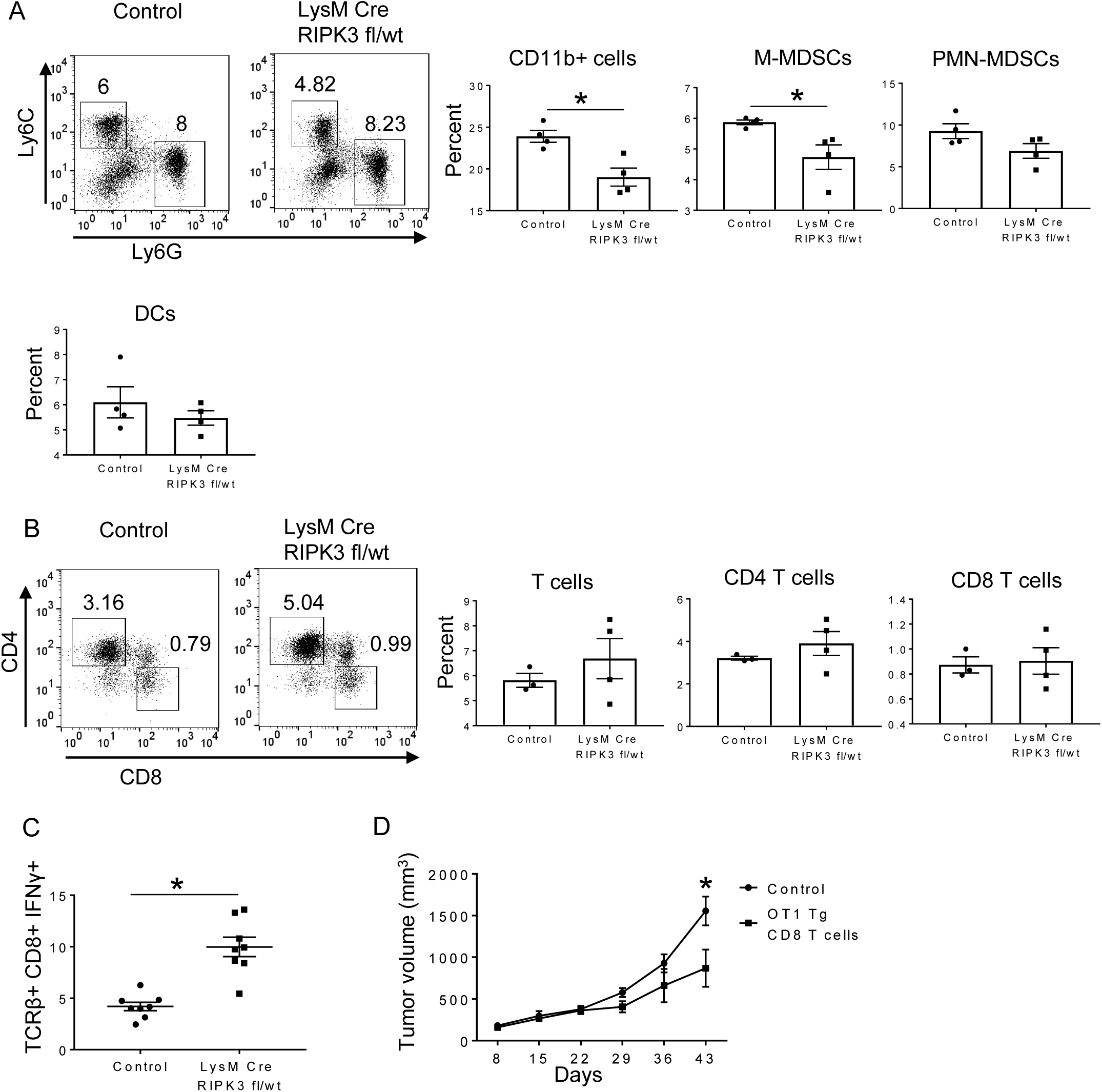
RIPK3 promotes suppression of CD8 T cells in KP lung tumor bearing mice. Total myeloid cells (CD11b+), M-MDSCs, PMN-MDSCs, and DCs (**A**), total T cells, CD4 and CD8 T cells (**B**) in WT and LysM Cre RIPK3 GFP fl/wt mice (n=4 mice/group). **C**. Splenic cells were stimulated with SIINFEKL peptide and CD8 T cells were analyzed for IFNγ. **D**. Mice with KP lung tumors were injected with OT1 Tg CD8 T cells that were in vitro stimulated with OVA antigen (n = 5 mice/group). * p< 0.05.

There was also some reduction in PMN-MDSCs and DCs. Overall, MDSCs including M-MDSCs and PMN-MDSCs were reduced in KP tumor bearing mice after partial deletion of RIPK3 in myeloid cells. T cells were slightly increased in LysM Cre RIPK3 fl/wt mice compared to WT mice; however, this was not significantly different (Fig. 2B). KP tumor cells express OVA antigen (12) and it is likely that OVA specific T cells are expanded in in LysM Cre RIPK3 fl/wt mice due to reduced MDSCs. To evaluate this rationale, splenic cells were activated with SIINFEKL peptide (OVA antigen) and T cells producing IFNγ were analyzed. CD8+ IFNγ+ T cells were increased in LysM Cre RIPK3 fl/wt mice compared to WT (Fig. 2C). This shows that reduction of RIPK3 in MDSCs generates more antigen specific CD8 T cells with cytotoxic activity, which could target KP tumor cells expressing OVA antigen. To show that antigen specific CD8 T cells directly inhibit KP tumor cells expressing OVA antigen, OT-1 Tg splenic cells (expressing TCR specific for OVA antigen SIINFEKL) were stimulated in vitro with SIINFEKL peptide to generate activated OT-1 CD8 T cells and adoptively transferred into KP tumor bearing mice (Fig. 2D). KP tumors were reduced in mice treated with OT-1 CD8 T cells compared to untreated mice showing that antigen specific CD8 T cells kill tumor cells expressing this antigen. Together, these results show that partial deletion of RIPK3 in myeloid cells including MDSCs improves the cytotoxic capacity of antigen specific CD8 T cells to inhibit KP tumor growth.

### Inhibition of RIPK3 with GSK 872 reduces KP lung tumors

The signaling function of RIPK3 can be inhibited by GSK 872, which was found to be highly specific for RIPK3 among several kinases (13). In this study, GSK 872 induces apoptosis at high concentrations in different cell lines such as epithelial cells, fibroblasts, and tumor cells. However, macrophages were viable for at least 18 hours when treated with high concentrations of GSK 872. Additionally, GSK 872 inhibited cytokine production in macrophages (10). Therefore, we reasoned that a relatively lower concentration of GSK 872 would inhibit signaling function and is unlikely to induce apoptosis. To evaluate the potency of GSK 872 in inhibiting KP lung tumor growth in vivo, mice with KP lung tumors between 50 and 100 mm^3^ were treated with GSK 872 every 3 days (Fig. 3A). Tumor growth was reduced compared to mice treated with vehicle. To assess if GSK 872 could inhibit tumor growth when tumors were larger, KP tumors between 200 to 300 mm^3^ were treated similarly (Fig. 3B). Treated mice had a reduction in tumor size compared to control mice. Although, this reduction in tumor volume was not significant, the inhibition of large tumors has translational significance because most NSCLC are diagnosed in the advanced stage (14). Therefore, GSK 872 could be used in combination with checkpoint inhibitors to effectively reduce tumor progression.

**Figure 3.**
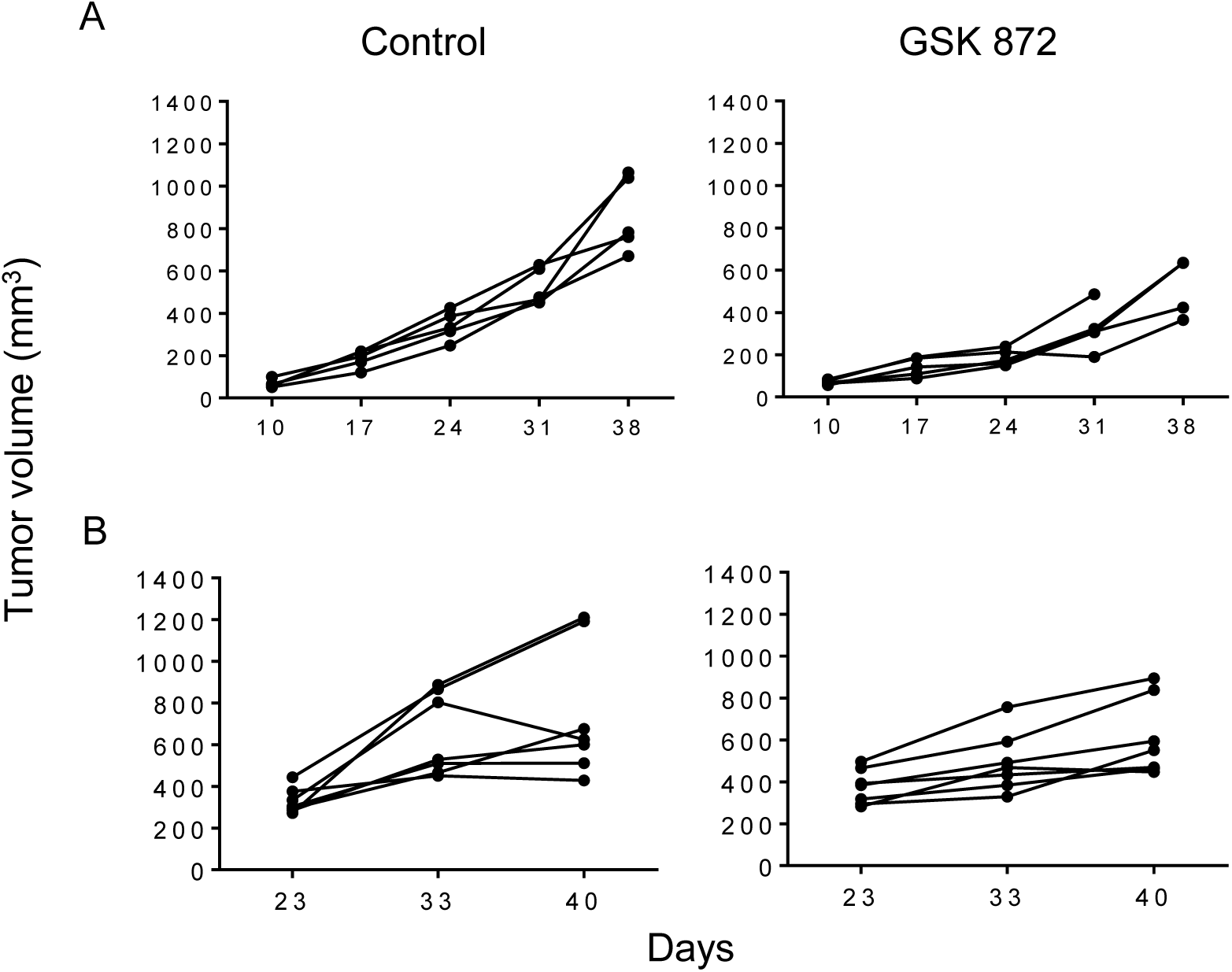
GSK 782 inhibits KP lung tumors. KP lung tumors were treated with RIPK3 inhibitor GSK 872 starting when tumors were between 50 and 100 mm^3^ (n=5/group) (**A**) and between 200 and 500 mm^3^ (n =7/ group) (**B**) for 4 weeks. p < 0.05 between control and GSK 872 treated group in A.

### Inhibition of RIPK3 increases the efficacy of checkpoint blockade

Efficacy of checkpoint blockade against NSCLC is reduced due to accumulation of regulatory cells including MDSCs. We show that deletion of RIPK3 in myeloid cells including MDSCs and a RIPK3 inhibitor GSK 872 reduces KP tumor growth. Therefore, GSK 872 could be used to target MDSCs that are abundant in cancer patients and are known to suppress antitumor T cells. To evaluate if GSK 872 could improve the efficacy of checkpoint inhibitors, KP tumors were treated with GSK 872 alone and GSK 872 with anti PD-1 or anti CTLA-4 antibodies. Tumor growth was reduced after treatment with GSK 872, anti PD-1, or anti CTLA-4 (Fig. 4A, B). Compared to single agent treatment, combination treatment with both GSK 872 and anti PD-1 or anti CTLA-4 reduced tumor growth more. This showed that GSK 872 could effectively reduce KP tumor growth by suppressing MDSC function in combination with checkpoint inhibitors to reduce exhaustion of CD8 T cells. We depleted CD8 T cells in mice treated with both GSK 872 and anti PD-1 or anti CTLA-4 to show that CD8 T cells are critical for inhibiting KP tumors. In the absence of CD8 T cells, KP tumor growth in combination treatment groups were similar to tumors in control mice (Fig. 4C). Therefore, GSK 872 with anti PD-1 or anti CTLA-4 improved the efficacy of CD8 T cells to inhibit the progression of KP tumors. GSK 872 likely inhibited the function of MDSCs to improve the efficacy of checkpoint blockade and is a valuable treatment option for treating NSCLC.

**Figure 4.**
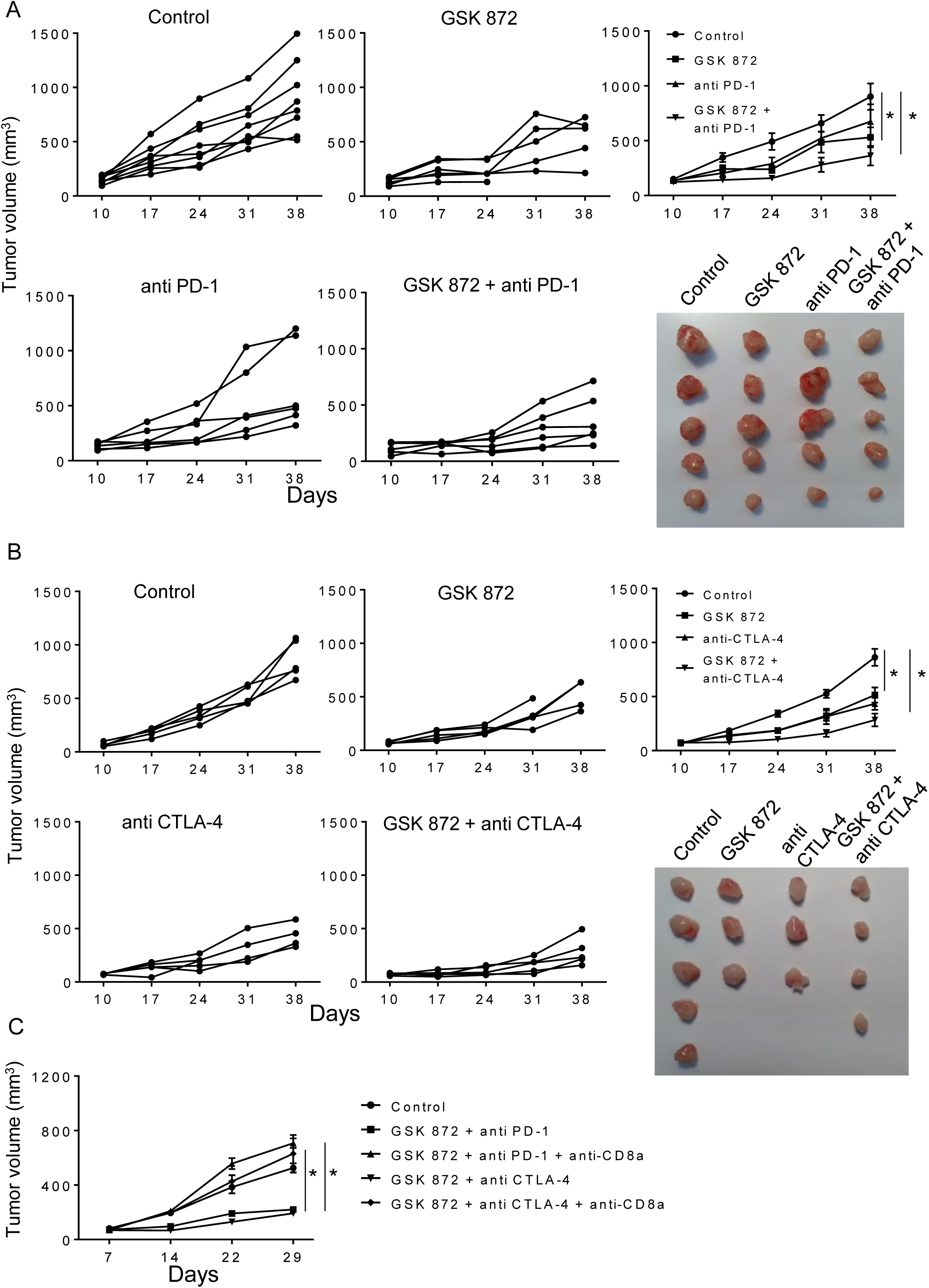
GSK 872 improves efficacy of checkpoint inhibitors. Mice with KP lung tumors were injected with GSK 872 with or without anti PD-1 (**A**) or anti CTLA-4 (**B**) every week. **C**. CD8 T cells were depleted with anti CD8 antibody in KP lung tumor bearing mice treated with GSK 872 with anti PD-1 or anti CTLA-4 (n = 5 to 8 mice/group). * p< 0.05.

M-MDSCs suppress both antigen specific and non-antigen specific T cells (7,15), whereas PMN-MDSCs target antigen specific CD8 T cells (16). Reduction of both M-MDSCs and PMN-MDSCs and tumor volume in tumor bearing mice with RIPK3 deletion in myeloid cells, suggests that the suppressive mechanism of MDSCs is inhibited. This is likely due to reduction of RIPK3 induced cytokines such as COX-2 and IL-1β in MDSCs (9). These cytokines promote the suppressive function of MDSCs (7). Therefore, deletion of RIPK3 in MDSCs would likely reduce the suppressive function of MDSCs and allow cytotoxic antigen specific CD8 T cells to inhibit tumor growth, which in turn results in fewer MDSCs. In NSCLC, more CD8 T cells expressing granzyme B and IFNγ results in better prognosis and survival (17). Consistently, targeting RIPK3 in KP tumor bearing mice generated more cytotoxic CD8 T cells. GSK 872, a specific inhibitor of RIPK3 inhibited both small and large KP tumors. This suggested that GSK 872 inhibited cytokine production in MDSCs and reduced their ability to suppress T cells. This is supported by our previous study, where GSK 872 inhibited cytokine production by MDSCs in vitro (9). The ability of GSK 872 to inhibit large KP tumors is particularly relevant as a potential addition in immunotherapeutic treatment for lung cancer. This is an effective strategy as shown by the improved efficacy of anti PD-1 and anti CTLA-4 in reducing KP lung tumor growth. Treatment of KP NSCLC is a challenge considering the dearth of therapies for NSCLC with KRAS mutation (18). GSK 872 is a viable alternative to existing treatments due to its ability to target RIPK3 mediated inflammatory cytokines and its tolerability in vivo.

---

**Supplementary figure 1.**
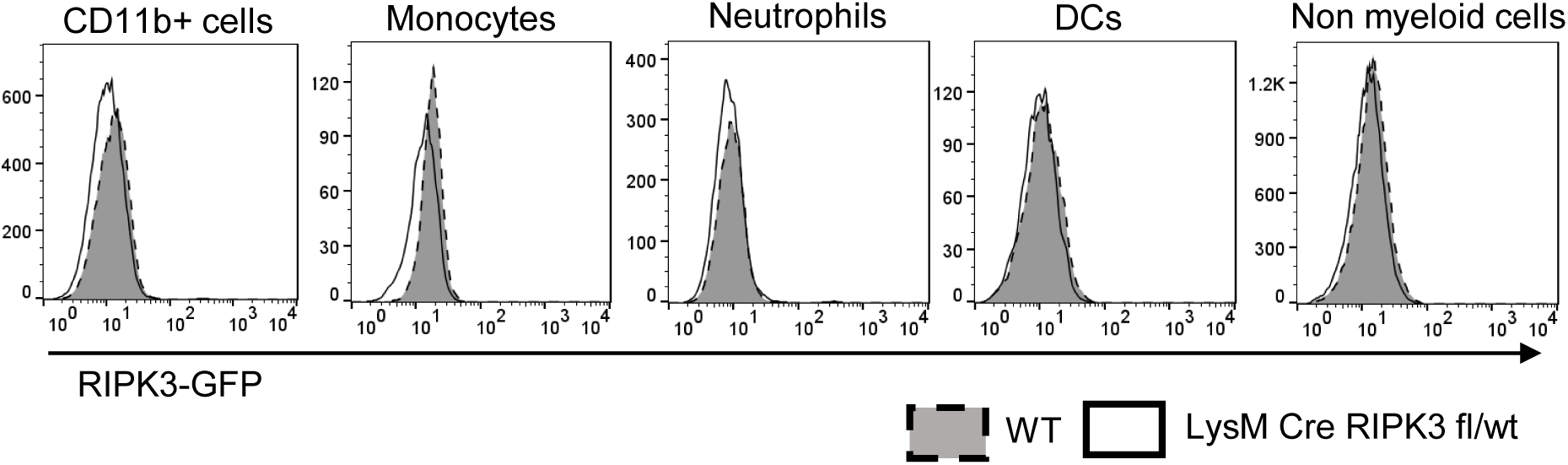
Deletion of RIPK3 in myeloid cells. RIPK3 – GFP floxed mice were bred with LysM Cre mice to generate LysM Cre RIPK3-GFP floxed/wt mice, which expressed reduced levels of RIPK3-GFP in splenic CD11b+ cells and monocytes.

**Supplementary figure 2.**
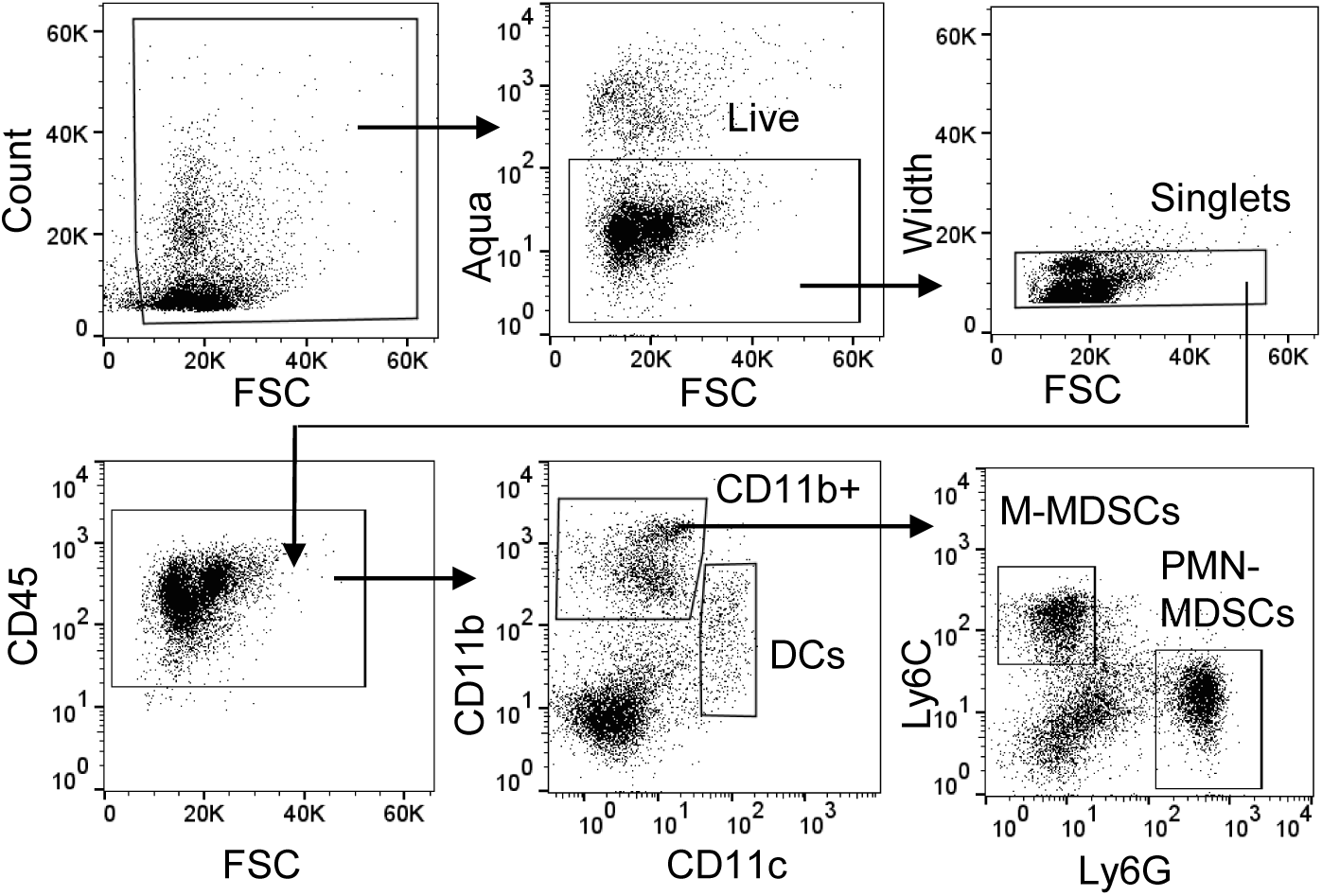
Gating strategy for flow cytometry analysis of splenocytes. Viable CD45+ mouse splenocytes were separated into CD11b+ cells and CD11c+ DCs. CD11b+ cells were separated into M-MDSCs (CD11b+Ly6C+Ly6Glo) and PMN-MDSCs (CD11b+Ly6G+LyC-).

## References

1. Bray F, Ferlay J, Soerjomataram I, Siegel RL, Torre LA, Jemal A. Global cancer statistics 2018: GLOBOCAN estimates of incidence and mortality worldwide for 36 cancers in 185 countries. CA Cancer J Clin 2018;68(6):394–424.

2. Ferrer I, Zugazagoitia J, Herbertz S, John W, Paz-Ares L, Schmid-Bindert G. KRAS-Mutant non-small cell lung cancer: From biology to therapy. Lung Cancer 2018;124:53–64.

3. Herbst RS, Morgensztern D, Boshoff C. The biology and management of non-small cell lung cancer. Nature 2018;553(7689):446–54.

4. Baumeister SH, Freeman GJ, Dranoff G, Sharpe AH. Coinhibitory Pathways in Immunotherapy for Cancer. Annu Rev Immunol 2016;34:539–73.

5. Trujillo JA, Sweis RF, Bao R, Luke JJ. T Cell-Inflamed versus Non-T Cell-Inflamed Tumors: A Conceptual Framework for Cancer Immunotherapy Drug Development and Combination Therapy Selection. Cancer Immunol Res 2018;6(9):990–1000.

6. Popovic A, Jaffee EM, Zaidi N. Emerging strategies for combination checkpoint modulators in cancer immunotherapy. J Clin Invest 2018;128(8):3209–18.

7. Marvel D, Gabrilovich DI. Myeloid-derived suppressor cells in the tumor microenvironment: expect the unexpected. J Clin Invest 2015;125(9):3356–64.

8. Wegner KW, Saleh D, Degterev A. Complex Pathologic Roles of RIPK1 and RIPK3: Moving Beyond Necroptosis. Trends Pharmacol Sci 2017;38(3):202–25.

9. Jayakumar A, Bothwell ALM. RIPK3-Induced Inflammation by I-MDSCs Promotes Intestinal Tumors. Cancer Res 2019;79(7):1587–99.

10. Najjar M, Saleh D, Zelic M, Nogusa S, Shah S, Tai A, et al. RIPK1 and RIPK3 Kinases Promote Cell-Death-Independent Inflammation by Toll-like Receptor 4. Immunity 2016;45(1):46–59.

11. Moriwaki K, Balaji S, Bertin J, Gough PJ, Chan FK. Distinct Kinase-Independent Role of RIPK3 in CD11c+ Mononuclear Phagocytes in Cytokine-Induced Tissue Repair. Cell Rep 2017;18(10):2441–51.

12. DuPage M, Cheung AF, Mazumdar C, Winslow MM, Bronson R, Schmidt LM, et al. Endogenous T cell responses to antigens expressed in lung adenocarcinomas delay malignant tumor progression. Cancer Cell 2011;19(1):72–85.

13. Mandal P, Berger SB, Pillay S, Moriwaki K, Huang C, Guo H, et al. RIP3 induces apoptosis independent of pronecrotic kinase activity. Mol Cell 2014;56(4):481–95.

14. How to detect lung cancer. Lung Cancer Foundation of America. Accessed 2019.

15. Gallina G, Dolcetti L, Serafini P, De Santo C, Marigo I, Colombo MP, et al. Tumors induce a subset of inflammatory monocytes with immunosuppressive activity on CD8+ T cells. J Clin Invest 2006;116(10):2777–90.

16. Youn JI, Nagaraj S, Collazo M, Gabrilovich DI. Subsets of myeloid-derived suppressor cells in tumor-bearing mice. J Immunol 2008;181(8):5791–802.

17. Gettinger SN, Choi J, Mani N, Sanmamed MF, Datar I, Sowell R, et al. A dormant TIL phenotype defines non-small cell lung carcinomas sensitive to immune checkpoint blockers. Nat Commun 2018;9(1):3196.

18. Pakkala S, Ramalingam SS. Personalized therapy for lung cancer: striking a moving target. JCI Insight 2018;3(15).

